# Fine-grained temporal mapping of derived high-frequency variants supports the mosaic nature of the evolution of *Homo sapiens*

**DOI:** 10.1101/2021.01.22.427608

**Authors:** Alejandro Andirkó, Juan Moriano, Alessandro Vitriolo, Martin Kuhlwilm, Giuseppe Testa, Cedric Boeckx

## Abstract

As our knowledge about the history of the *Homo sapiens* lineage becomes increasingly complex, large-scale estimations of the time of emergence of derived variants become essential to be able to offer more precise answers to time-sensitive hypotheses concerning human evolution. Using an open repository of genetic variant age estimations recently made available, we offer here a temporal evaluation of various evolutionarily relevant datasets, such as *Homo sapiens*-specific variants, high-frequency variants found in genetic windows under positive selection, introgressed variants from extinct human species, as well as putative regulatory variants in various brain regions. We find a recurrent bimodal distribution of high-frequency variants, but also evidence for specific enrichments of gene categories in various time windows, which brings into prominence the 300-500k time slice. We also find evidence for very early mutations impacting the facial phenotype, and much more recent molecular events linked to specific brain regions such as the cerebellum or the precuneus. Additionally, we present a case study of an evolutionarily relevant gene, *BAZ1B*, and its targets, to emphasize the importance of applying temporal data to specific evolutionary questions. Overall, we present a unique resource that informs and complements our previous knowledge of *Homo sapiens* evolution using publicly available data, and reinforce the case for the mosaic, temporally very extended nature of the evolutionary trajectory of our species.

## 1 Introduction

The past decade has seen a significant shift in our understanding of the evolution of our lineage. We now recognize that anatomical features used as diagnostic for our species (globular neurocranium, small, retracted face, presence of a chin, narrow trunk, to cite only a few of the most salient traits associated with ‘anatomical modernity’) did not emerge as a package, from a single geographical location, but rather emerged gradually, in a mosaic-like fashion across the entire African continent [1]. Likewise, behavioral characteristics once thought to be exclusive of *Homo sapiens* (funerary rituals, parietal art, ‘symbolic’ artefacts, etc.) have recently been attested in some form in closely related (extinct) clades, casting doubt on a simple definition of ‘cognitive/behavioral’ modernity [2]. We have also come to appreciate the extent of (multidirectional) gene flow between Sapiens and Neanderthals and Denisovans, raising interesting questions about speciation [3, 4, 5, 6]. Last, but not least, it is now well established that our species has a long history. Robust genetic analyses [7] indicate a divergence time between us and other hominins for which genomes are available of roughly 700kya, leaving perhaps as many as 500ky between then and the earliest fossils displaying a near-complete suite of modern traits (Omo Kibish 1, Herto 1 and 2) [8].

Such a long period of time allows for the distinction between early and late members of our species [8]. Genomic analysis of ancient human remains in Africa reveal deep population splits and complex admixture patterns among populations well before the coalescence of modernity in the fossil record [9, 10]. At the same time, reanalysis of archaic fossils in Africa [11] point to the extended presence of multiple hominins on this continent, with the possibility of ‘super-archaic’ admixture [12, 13]. Lastly, our deeper understanding of other hominins point to derived characteristics in these lineages that make some of our species’ traits more ancestral (less ‘modern’) than previously believed [14].

In the context of this significant rewriting of our history, we decided to explore the temporal structure of an extended catalog of single nucleotide changes found at high frequency (HF*≥* 90%) across major modern populations we previously generated on the basis of 3 high-coverage archaic genomes [15]. This catalog aims to offer a richer picture of molecular events setting us apart from our closest extinct relatives. To do so, we took advantage of the Genealogical Estimation of Variant Age (GEVA) tool [16]. GEVA is a coalescence-based method that provides age estimates for over 45 million human variants. GEVA is non-parametric, making no assumptions about demographic history, tree shapes, or selection. (For additional details on GEVA, see section 4). Our overall objective here is to use the temporal resolution afforded by GEVA to to estimate the age of emergence of polymorphic sites, and gain further insights into the complex evolutionary trajectory.

Here, we reveal a bimodal temporal distribution of modern human derived high-frequency variants and provide insights into milestones of *Homo sapiens* evolution through the investigation of the molecular correlates and the predicted impact of variants across evolutionary-relevant periods. Our chronological atlas allows us to provide a time window estimate of introgression events and evaluate the age of variants associated with signals of positive selection, as well as estimate the age of enhancer regulatory variants for different brain regions. Our enrichment analyses uncovers GO-terms unique to specific temporal windows, prominently facial and behavioral-related terms between 300k and 500k years. With a finer-grained level of scrutiny, our machine learning-based analyses predicting differential gene expression regulation of mapped variants (through [17]) reveals a trend towards downregulation in the aforementioned period (300k-500k years; corresponding to the early emergence of our species). We further identify variant-associated genes whose differential regulation may specifically affect brain structures thought to be derived in late *Homo sapiens* such as the cerebellum and the precuneus. Finally, we delved into the study of *BAZ1B*, for its contribution to our understanding of craniofacial development and human evolution [18]. We found a cluster of variants linked to a specific set of *BAZ1B* targets dated around 300-500k years (within the suggested period of appearance of distinctive facial traits in our species), and characterized a set of older variants that further shed light into the timing of the emergence of the ‘modern’ human face.

## 2 Results

The distribution of alleles over time follows a bimodal distribution regardless of the frequency cutoff (Figure 1A; Figure S1), with a global maximum around 40kya (for complete allele counts, see section 4). The two modes of the distribution correspond to two periods of significance in the evolutionary history of *Homo sapiens*. The more recent peak of HF variants arguably corresponds to the period of population dispersal around 100kya [19], while the older distribution contains the period associated with the divergence between *Homo sapiens* and other *Homo* species [7, 20]. When dividing the modes (at the 300kya time mark), the distribution of variants over time is statistically different between the set of overall derived variants and each of the two HF filtered sets (*p <* 0.01, Kolmogorov–Smirnov test).

**Figure 1.**
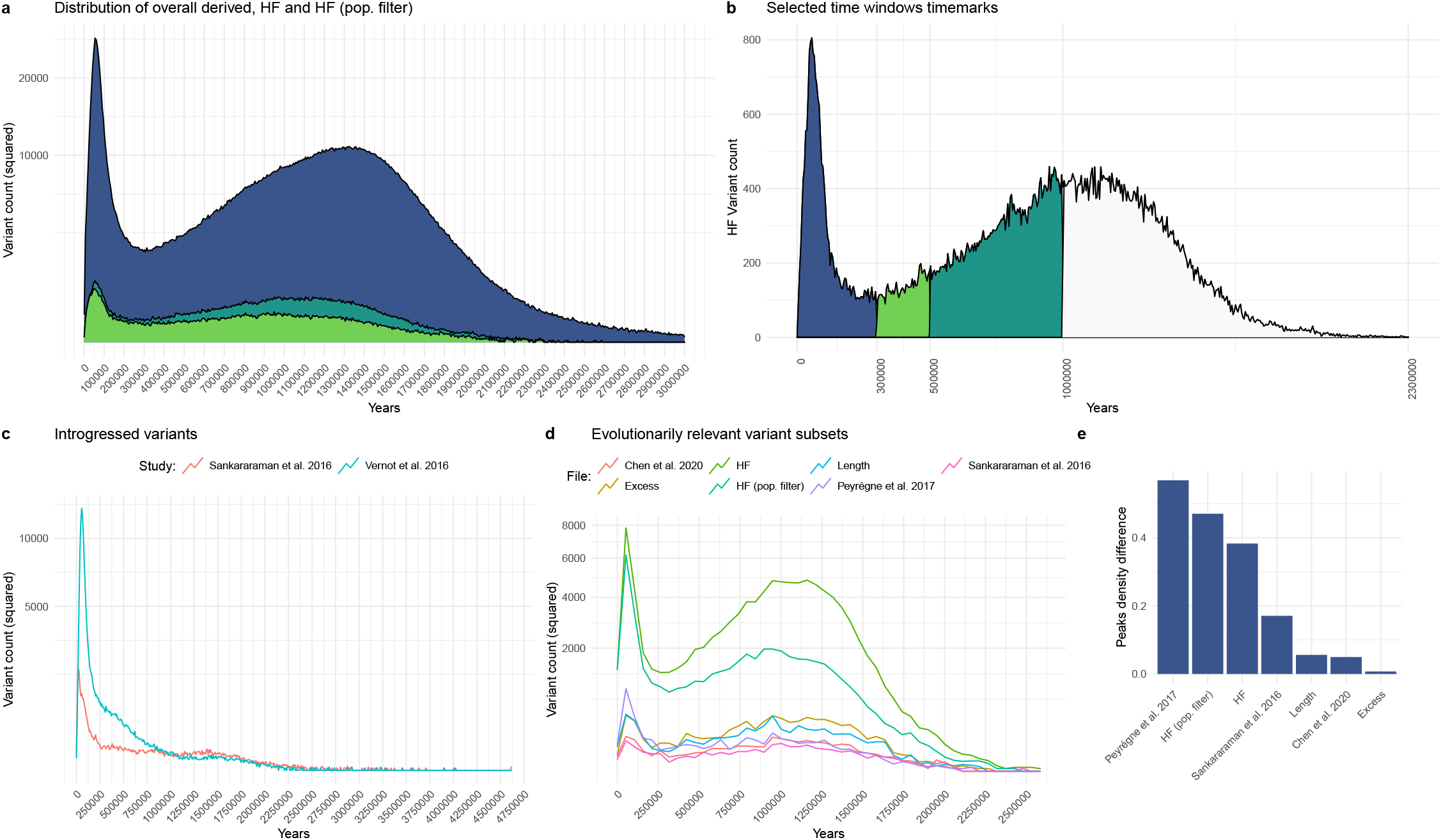
A: Distribution of derived *Homo sapiens* alleles over time with no frequency cutoff, in HF and the modified population-wise HF subset (see sec. 4). Trimmed at 3mya – the full distributions is shown in Fig S1 B: Selected chronological milestones used in our study, as informed by the archaeological record. C: Distribution of introgressed alleles over time, as identified by [23] and [26]. D: Plots of HF variants in datasets relevant to human evolution, including regions under positive selection [25], regions depleted of archaic introgression [23, 24] and genes showing an excess of HF variants (‘excess’ and ‘length’) [15]. Variant counts in A, C and D are squared to aid visualization. E: Kernel density difference between the highest point in the distributions of D (leftmost peak) and the second, older highest density peak, normalized, in percentage units.

In order to divide the data for downstream analysis we considered a *k*-means clustering analysis (at *k* = 3 and *k* = 4, Figure S2). This clustering method yields a division clear enough to distinguish between early and late *Homo sapiens* specimens after the split with other human species. However, we reasoned that such a k-means division is not precise enough to represent key milestones used to test specific time-sensitive hypotheses. For this reason, we adopted a literature-based approach, establishing different cutoffs adapted to the need of each analysis below (Figure 1B). Our basic division consisted of three periods: a recent period from the present to 300 thousand years ago (kya), the local minimum, roughly corresponding to the period considered until recently to mark the emergence of *Homo sapiens*; a later period from 300kya to 500kya, the period associated with earlier members of our species such as the Jebel Irhoud fossil [21] ; and a third, older period, from 500kya to 1 million year ago, corresponding to the time of the most recent common ancestor with the Neanderthal and Denisovan lineage [22]. Finer-grained time slices were adopted for further analyses (see, e.g., section 2.3).

We note that the distribution goes as far back as 2.5 million years ago (see Figure 1A) in the case of HF variants, and even further back in the case of the derived variants with no HF cutoff. This could be due to our temporal prediction model choice (GEVA clock model, of which GEVA offers three options, as detailed in 4), as changes over time in human recombination rates might affect the timing of older variants [16], or to the fact that we don’t have genomes for older *Homo* species. Some of these very old variants may have been inherited from them, and lost further down the archaic lineages. In this context, we note that 40% of the genes that exhibit an excess of mutations in the modern lineage and totally lack HF derived variants in other hominins in [15] do not exhibit any single ‘recent’ (<400kya) HF variant (Fig. S3).

### 2.1 Variant subset distributions

In an attempt to see if specific subsets of variants had strikingly different distributions over time, we selected a series of evolutionary relevant sets of data publicly available, such as genome regions depleted of archaic introgression (so-called ‘deserts of introgression’) [23, 24], and regions under putative positive selection [25], and mapped the HF variants from [15] falling within those regions. We also examined genes that accumulate more HF variants than expected given their length and in comparison to the number of mutations these genes accumulate on the archaic lineages (‘length’ and ‘excess’ lists from [15] – see sec. 4). Finally, we plotted introgressed alleles [23, 26]. A bimodal distribution is clearly visible in all the subsets except the introgression datasets (Figure 1C). Introgressed variants peak locally in the earlier period (0-100kya). The distribution roughly fades after 250kya, in consonance with the possible timing of introgression events [4, 12, 24, 27]. As a case example, we plotted those introgressed variants associated with phenotypes highlighted in Table 1 of [28]. As shown in Figure S4, half of the variants cluster around the highest peak, but other variants may have been introduced in earlier instances of gene flow. We caution, though, that multiple (likely) factors, such as gene flow from Eurasians into Africa, or effects of positive selection affecting frequency, influence the distribution of age estimates and make it hard to draw any firm conclusions. We also note that the two introgressed variant counts, derived from the data of [26] and [23], follow a significantly different distribution over time (*p <* 2.2 *−* 16, Kolmogorov–Smirnov test) (Figure 1C).

**Table 1.**
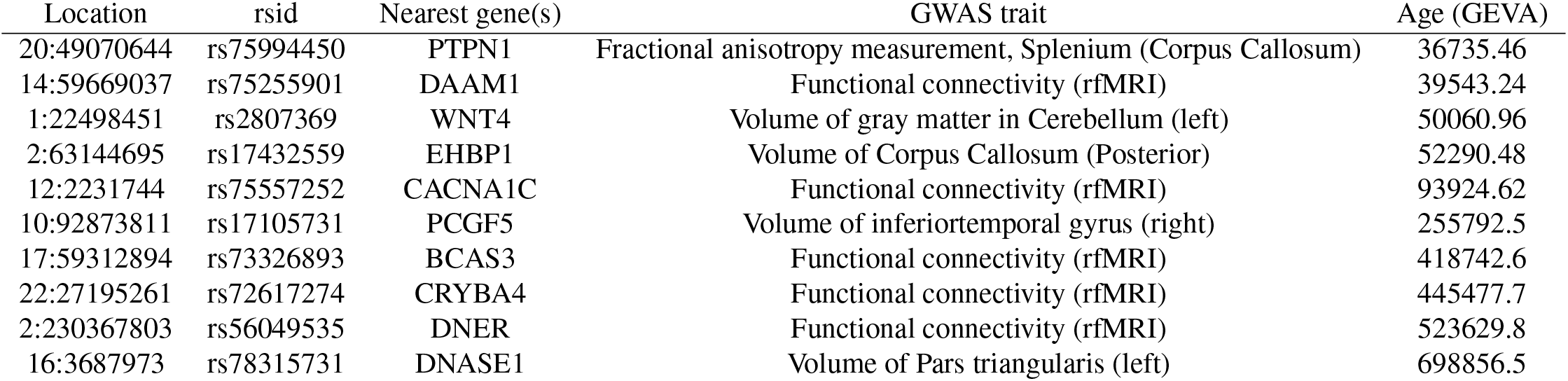
Big40 Brain volume GWAS [41] top hits with high predicted gene expression in ExPecto (*log >* 0.01, RPKM), along with dating as provided by *GEVA*. ‘Functional connectivity’ is a measure of temporal activity synchronization between brain parcels at rest (originally defined in [46]).

Finally, we examined the distribution of putatively introgressed variants across populations, focusing on low-frequency variants whose distributions vary when we look at African vs. non-African populations (Figure S5). As expected, those variants that are more common in non-African populations are found in higher proportions in both of the Neanderthal genomes studied here, with a slightly higher proportion for the Vindija genome, which is in fact assumed to be closer to the main source population of introgression. We detect a smaller contribution of Denisovan variants overall, which is expected on several grounds: given the likely more frequent interactions between modern humans and Neanderthals, the Denisovan individual whose genome we relied on is likely part of a more pronounced “outgroup”. Gene flow from modern humans into Neanderthals also likely contributed to this pattern.

In the case of the regions under putative positive selection, we find that the distribution of variant counts has a local peak in the most recent period (0-100kya) that is absent from the deserts of introgression datasets. Also, as shown in 1E, the distribution of variant counts in these regions under selection shows the greatest difference between the two peaks of the bimodal distribution. Still, we should stress that our focus here is on HF variants, and that of course not all HF variants falling in selective sweep regions were actual targets of selection. Figure S6 illustrates this point for two genes that have figured prominently in early discussions of selective sweeps since [3]: *RUNX2* and *GLI3*. While recent HF variants are associated with positive selection signals (indicated in purple), older variants exhibit such associations as well. Indeed some of these targets may fall below the 90% cutoff chosen in [15]. In addition, we are aware that variants enter the genome at one stage and are likely selected for at a (much) later stage [29, 30]. As such our study differs from the chronological atlas of natural selection in our species presented in [31] (as well as from other studies focusing on more recent periods of our evolutionary history, such as [32]). This may explain some important discrepancies between the overall temporal profile of genes highlighted in [31] and the distribution of HF variants for these genes in our data (Figure S7).

Having said this, our analysis recaptures earlier observations about prominent selected variants, located around the most recent peak, concerning genes such as *CADPS2* ([33], Fig. S8). This study also identifies a large set of old variants, well before 300kya, associated with genes belonging to putative positively-selected regions before the deepest divergence of *Homo sapiens* populations [34], such as *LPHN3, FBXW7*, and *COG5* (figure S9).

Finally, we estimated the age of putative regulatory variants of the prefrontal (PFC), temporal (TC) and cerebellar cortices (CBC), using the large scale characterization of regulatory elements of the human brain provided by the PsychENCODE Consortium [35]. We did the same for the modern human HF missense mutations [15]. A comparative plot reveals a similar pattern between the three structures, with no obvious differences in variant distribution (see Fig. S10). The cerebellum contains a slightly higher number of variants assigned to the more recent peak when the proportion to total mapped variants is computed: 15.59% to 14.97% (PFC) and 15.20% (TC). We also note that the difference of dated variants between the two local maxima is more pronounced in the case of the cerebellum than in the case of the two cortical tissues, whereas this difference is more reduced in the case of missense variants (Fig. S10).We caution, though, that the overall number of missense variants is considerably lower in comparison to the other three datasets.

### 2.2 Gene Ontology analysis across temporal windows

In order to interpret functionally the distribution of HF variants in time, we performed enrichment analyses accessing curated databases via the *gProfiler2* R package [36]. For the three time windows analyzed (corresponding to the recent peak: 0-300kya; divergence time and earlier peak: 500kya-1mya; and time slot between them: 300kya-500kya), we identified unique and shared gene ontology terms (see Figure 2A and sec. 4). Of note, when we compared the most recent period against the two earlier windows together (from 300kya-1mya), we found bone, cartilage and visual system-related terms only in the earlier periods (hypergeometric test; adj. *p <* 0.01; Table S1). Further differences are observed when thresholding by an adjusted *p <* 0.05. In particular, terms related to behavior (startle response), facial shape (narrow mouth) and hormone systems only appear in the middle (300-500k) period (Table S2; Figure S11). A summary of terms shared across the three time windows can be seen in Figure S12.

**Figure 2.**
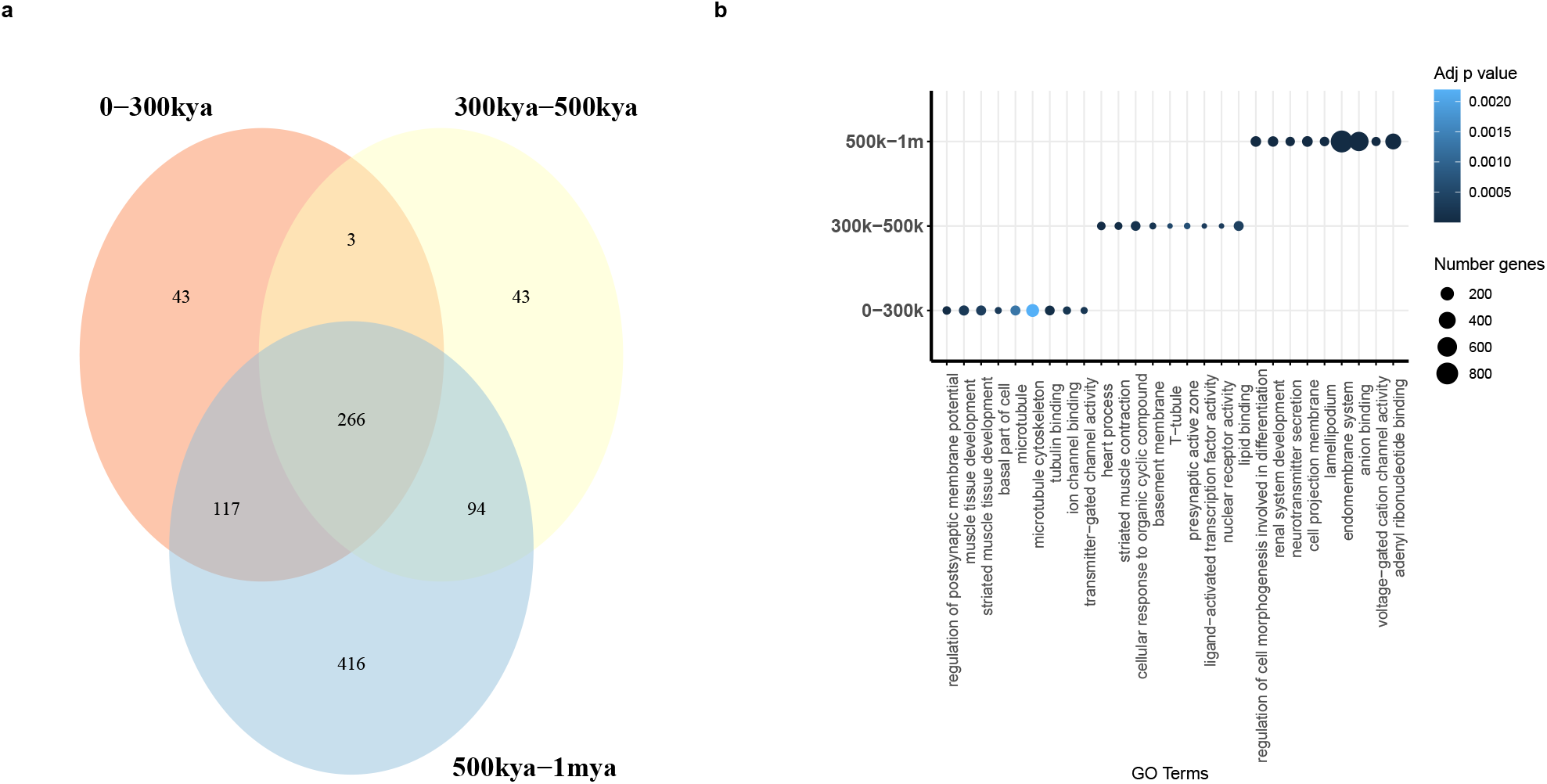
A: Venn diagram of GO terms associated with genes shared across time windows. B: Top GO terms per time window.

### 2.3 Gene expression predictions

To see if term-enriched genes are associated with particular expression profiles, we made use of ExPecto [17], a sequence-based tool to predict gene expression *in silico* (see description in section 4). We found that there is a significant skewness towards extreme negative values in the 300kya to 500kya time period that is not so salient in the other windows (as shown in quantile-quantile plots in Fig. S14). This skewness is present but not so salient in the overall set of tissue HF variant-specific expression predictions. A series of Kruskal-Wallis tests show that variants coming from GO-enriched genes have significant differences in their average expression levels in each period (0-300kya, 300-500kya and 500-800kya) compared to the others (*p* = 3.411*e −* 05, *p* = 4.032*e −* 08 and *p* = 4.032*e −* 08, adjusted by Bonferroni).

We applied the ExPecto tool as well to the overall derived HF variant dataset derived from [15], with a particular focus on expression changes in brain tissues.

To examine if certain tissues had a specially high predicted expression value in certain key time windows, we further divided the variants in six chronological groups ranging from the present to an estimated 800kya according to the GEVA set dating (Fig. 3A – see Fig. S15 for full details). Of note is the presence of the cerebellum in a period preceding the last major Out-of-Africa event (as predicted by [37]) in a landscape otherwise dominated by tissues such as the Adrenal Gland, the Pituitary, Astrocytes, and Neural Progenitor Cells.

**Figure 3.**
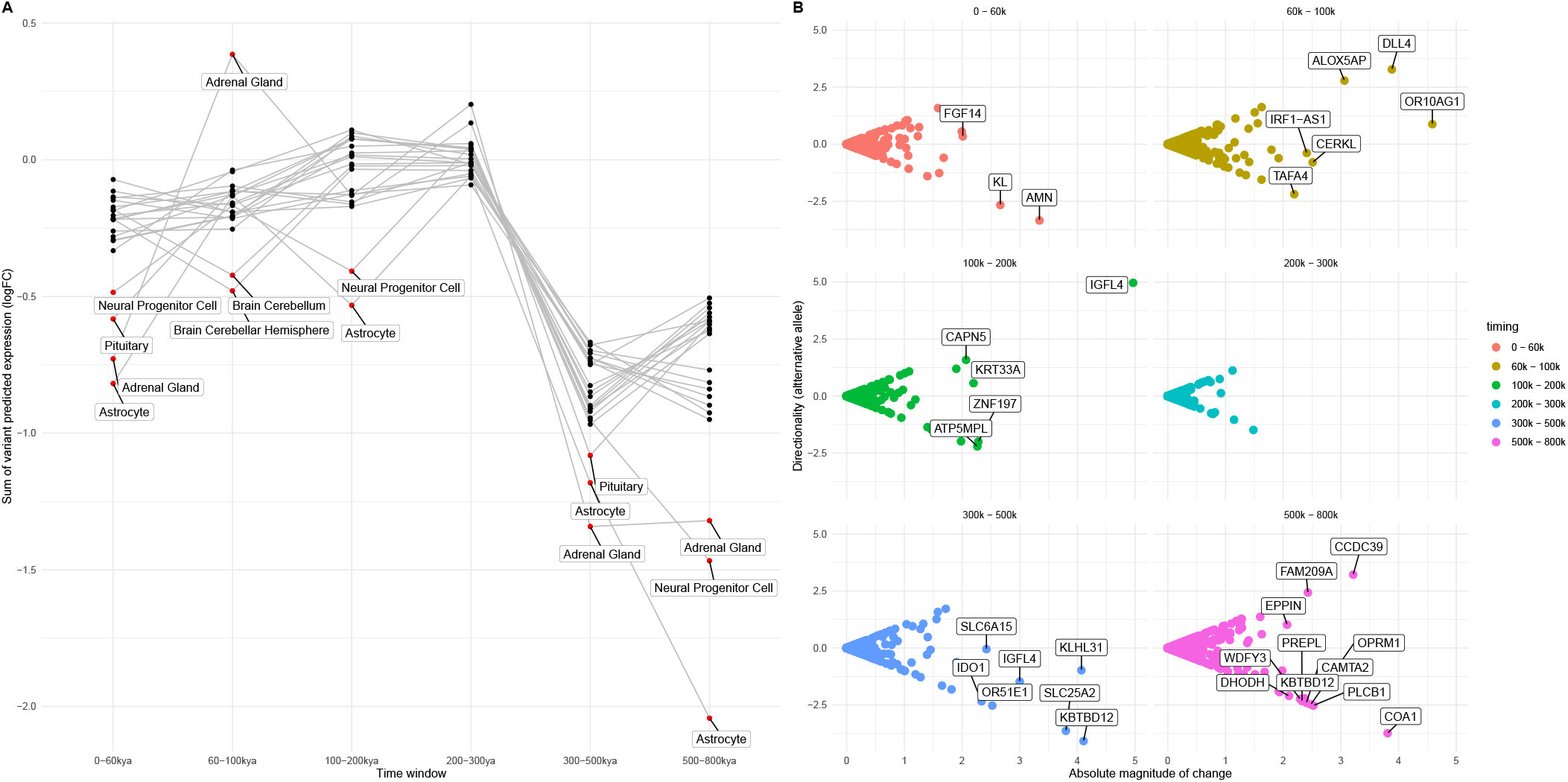
A: Sum of all directional mutation effects within 1kb to the TSS per time window in 22 brain regions from the ENCODE, GTEx and Road map datasets. Highlighted in red, bottom and top values labelled for illustration. Note, however, that expression values predicted are significantly different across time windows but not tissues (as detailed in sec. 2.3). B: Genes with a high sum of all directional mutation effects, and cumulative directionality of expression values.

The six windows (0-60, 60-100, 100-200, 200-300, 300-500 and 500-800kya) attempt to capture events in a finer-grained fashion (see sec. 4). We found that the sum of predicted gene expression values differs across timing windows, as determined by an approximate Kruskal-Wallis Test with random sampling (*n* = 1000) test, but not across tissues. A post-hoc Dunn test shows that expression values predicted by ExPecto are significantly different between the 60-100 and the 200-300 and 300-500 windows (*p* = 0.001 and *p* = 0.0012, p-values adjusted with Benjamini-Hochberg) and between 0-60 and 60-100 (*p* = 0.0102, adjusted). We performed an additional analysis to check whether there is an association between exact dates predicted by the GEVA tool and expression (as opposed to a time window division). The correlation between these two values is not significant (*p* = 0.3287, Pearson correlation test).

The authors of the article describing the ExPecto tool [17] suggest that genes with a high sum of absolute variant effects in specific time windows tend to be tissue or condition-specific. We explored our data to see if the genes with higher absolute variant effect were also phenotypically relevant (Figure 3B). Among these we find genes such as *DLL4*, a Notch ligand implicated in arterial formation [38]; *FGF14*, which regulates the intrinsic excitability of cerebellar Purkinje neurons [39]; *SLC6A15*, a gene that modulates stress vulnerability through the glutamate system [40]; and *OPRM1*, a modulator of the dopamine system that harbors a HF derived loss of stop codon variant in the genetic pool of modern humans but not in that of extinct human species [15].

We also crosschecked if any of the variants in our high-frequency dataset with a high predicted expression value (RPKM variant-specific values at *log >* 0.01) were found in GWASs related to brain volume. The Big40 UKBiobank GWAS meta-analysis [41] shows that some of these variants are indeed GWAS top hits and can be assigned a date (see Table 1). Of note are phenotypes associated with the posterior Corpus Callosum (Splenium), precuneus, and cerebellar volume. In addition, in a large genome-wide association meta-analysis of brain magnetic resonance imaging data from 51,665 individuals seeking to identify specific genetic loci that influence human cortical structure [42], one variant (rs75255901) in Table 1, linked to *DAAM1*, has been identified as a putative causal variant affecting the precuneus. All these brain structures have been independently argued to have undergone recent evolution in our lineage [37, 43, 44, 45], and their associated variants are dated amongst the most recent ones in the table.

### 2.4 Case study

As a case example of the potential of the GEVA dataset when applied to evolutionary questions, we examined HF variants found in *BAZ1B* and target genes. *BAZ1B* is a gene implicated in craniofacial defects in Williams-Beuren syndrome. We recently positioned this gene upstream in the developmental hierarchy of the modern human face on the basis of empirical evidence gathered from neural crest models with interfered gene function [18]. We wanted to determine if HF mutations harbored by *BAZ1B* are temporally accompanied by HF variant changes in a range of target genes that we previously demonstrated cluster in statistically significant ways when examined in an evolutionary context [18]. These targets fall in two broad groups: those genes whose expression patterns change in the same direction as that of *BAZ1B* (labeled “DIR”), and those whose expression patterns go in the opposite direction (labeled “INV”). Experimental validation further refined these two sets of genes and identified *bona fide* direct targets of *BAZ1B* (27DIR and 25INV genes, and, with further filtering, 13DIR and 17INV). We already observed that these two sets of targets overlap significantly with genes harboring (regulatory) HF mutations in modern human genomes compared to archaic human genomes, although for the broadest set of “INV” targets, the overlap resulted statistically significant for extinct human species as well [18].

Out of a total of 289 HF mutations harbored by direct targets of *BAZ1B*, 238 could be mapped via GEVA (Figure 4A-B). We observe that close to 25% of all HF variants associated with both INV and DIR targets are found in the oldest time slices defined by the occurrence of *BAZ1B* HF variants, around 1.3mya. 13% of all these ‘target’ variants are found in the 300-500k time window, and about the same percentage (15%) in the most recent (0-300k) period. In other words, unlike the general variant distribution found throughout this study, we do not find a recent peak of variants associated with *BAZ1B* targets. This is in line with the GO-enrichment results presented above, where we don’t find any enrichment for ‘face’-related terms in the most recent periods.

**Figure 4.**
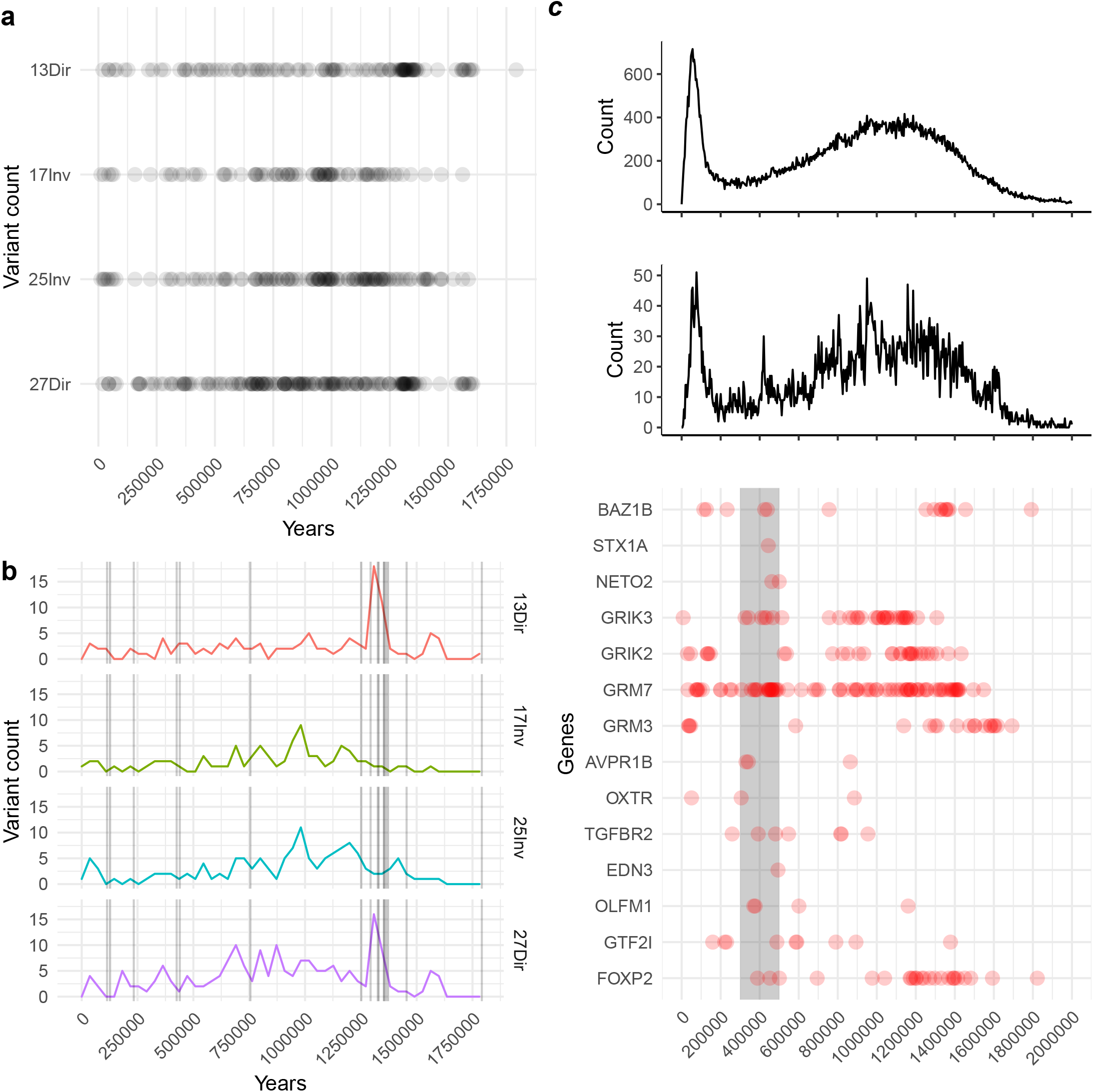
A: Accumulation of variants over time in genes whose expression levels are robustly correlated, directly (‘Dir’) or inversely (‘Inv’), with BAZ1B expression, as per [18]. B: Relation of variant emergence and BAZ1B mutations (vertical black lines) per list of robustly correlated target genes. C: Distribution of HF variants (top), variants in genes showing an excess of HF mutations (middle), and date of emergence of HF variants in selected genes over time (bottom), including a highlight between 300kya and 500kya (in gray). The total number of mapped HF variants for these genes follows a linear relationship with gene length (Fig. S. 18).

These results invited us to look more closely into the 300-500k period, which as been independently linked to the emergence of modern facial traits (Jebel Irhoud fossil, [21]), and possibly mark a change in our prosociality captured by the “self-domestication hypothesis” ([47, 48]). This period shows a local increase in HF variants for genes harboring an “excess” of mutations compared to archaics, controlling for gene length [15] (Fig 4C). Mutations in other genes we have previously linked to the earliest stages of self-domestication [49] cluster around this period, as shown in Fig 4C. Among them are other genes belonging to the Williams-Beuren Syndrome critical region (*STX1A, GTF2I*), prominent targets of *BAZ1B* implicated in Neural Crest processes (*OLFM1, EDN3, TGFBR2*), as well as specific classes of genes that modulate glutamate signaling (*GRIK3, GRIK2, GRM7, NETO2*) and hormones (*OXTR, AVPR1B*). Interestingly, the most recent HF variants in *FOXP2* we could map belong to that period.

It is noteworthy that HF variants harbored by genes associated with face and vocal tract anatomy that were singled out for their extensive methylation changes in [50] (*SOX9, ACAN, COL2A1, NFIX* and *XYLT1*) cluster (together with other *BAZ1B* HF mutations) in our dataset in a more recent time window (Fig S16), pointing to further refinement of the modern facial phenotype, in line with the authors’ own claims in [50]. It is also worth pointing out that *BAZ1B* (and its targets) harbor several HF mutations going back to as early as 900k, which may indicate that aspects of the ‘modern’ face are indeed as old as some have recently claimed, relying on a characterization of both proteomic and phenotypic characterizations of *Homo antecessor* [14, 51].

## 3 Discussion

Deploying GEVA to probe the temporal structure of the extended catalog of HF variants distinguishing modern humans from their closest extinct relatives ultimately aims to contribute to the goals of the emerging attempts to construct a molecular archaeology [52] and as detailed a map as possible of the evolutionary history of our species. Like any other archaeology dataset, ours is necessarily fragmentary. In particular, fully fixed mutations, which have featured prominently in early attempts to identify candidates with important functional consequences [52], fell outside the scope of this study, as GEVA can only determine the age of polymorphic mutations in the present-day human population. By contrast, the mapping of HF variants was reasonably good, and allowed us to provide complementary evidence for claims regarding important stages in the evolution of our lineage. This in and of itself reinforces the rationale of paying close attention to an extended catalog of HF variants, as argued in [15].

While we wait for more genomes from more diverse regions of the planet and from a wider range of time points, we find our results encouraging: even in the absence of genomes from the deep past of our species in Africa, we were able to provide evidence for different epochs and classes of variants that define these. Indeed, the emerging picture is very much mosaic-like in its character, in consonance with recent work in archeology [1].

Our analysis highlights the importance of a temporal window between 300-500k that may well correspond to a significant behavioral shift in our lineage, corresponding to the Jebel Irhoud fossil, but also in other parts of the African continent, to increased ecological resource variability [53], and evidence of long-distance stone transport and pigment use [54]. Other aspects of our cognitive and anatomical modernity emerged much more recently, in the last 150000 years, and for these our analysis points to the relevance of gene expression regulation differences in recent human evolution, in line with [55, 56, 57]. These two salient temporal windows are well represented by the density of HF mutations in genes such as *PTEN*, one of the genes highlighted in [15] as harboring an excess of derived HF mutations on the modern compared to extinct human lineages (Fig S17).

Lastly, our attempt to date the emergence of mutations in our genomes points to multiple episodes of introgression, whose history is likely to turn out to be quite complex.

## 4 Methods

### Homo sapiens variant catalog

We made use of a publicly available dataset [15] that takes advantage of the Neanderthal and Denisovan genomes to compile a genome-wide catalog of *Homo sapiens*-specific variation (genome version *hg19*, 1000 genomes project frequency data, dbSNP database). In addition to the full data, the authors offered a subset of the data that includes derived variants at a *≥* 90% global frequency cutoff. Since such a cutoff allows some variants to reach less than 90% in certain populations, as long as the total is *≥* 90%, we also considered including a metapopulation-wide variant *≥* 90% frequency cutoff dataset to this study (Fig 1A). All files (the original full and high-frequency sets and the modified, stricter high-frequency one) are provided in the accompanying code.

### GEVA

The Genealogical Estimation of Variant Age (GEVA) tool [16] uses a hidden Markov model approach to infer the location of ancestral haplotypes relative to a given variant. It then infers time to the most recent ancestor in multiple pairwise comparisons by coalescent-based clock models. The resulting pairwise information is combined in a posterior probability measure of variant age. We extracted dating information for the alleles of our dataset from the bulk summary information of GEVA age predictions. The GEVA tool provides several clock models and measures for variant age. We chose the mean age measure from the joint clock model, that combines recombination and mutation estimates. While the GEVA dataset provides data for 1000 genomes project and the Simons Genome Diversity Project, we chose to extract only those variants that were present in both datasets. Ensuring a variant is present in both databases implicitly increases genealogical estimates (as detailed in Supplementary document 3 of [16]), although it decreases the amount of sites that can be looked at. We give estimated dates after assuming 29 years per generation, as suggested in [58]. While other measures can be chosen, this value should not affect the nature of the variant age distribution nor our conclusions.

Out of a total of 4437804 for our total set of variants, 2294023 where mapped in the GEVA dataset (51% of the original total). For the HF subsets, the mapping improves: 101417 (74% of total) and 48424 (69%) variants were mapped for the original high frequency subset and the stricter, meta-population cutoff version, respectively.

### ExPecto

In order to predict gene expression we made use of the *ExPecto* tool [17]. *ExPecto* is a deep convolutional network framework that predicts tissue-specific gene expression directly from genetic sequences. *ExPecto* is trained on histone mark, transcription factor and DNA accessibility profiles, allowing *ab initio* prediction that does not rely on variant information training. Sequence-based approaches, such as the one used by *Expecto*, allow to predict the expression of high-frequency and rare alleles without the biases that other frameworks based on variant information might introduce. We introduced the high-frequency dated variants as input for *ExPecto* expression prediction, using the default tissue training models trained on the GTEx, Roadmap genomics and ENCODE tissue expression profiles. We then selected brain and brain-related tissues (as detailed in the code), and divided the variants by time period (0-60kya, 60-100kya, 100-200kya, 200-300kya, 300-500kya and 500-800kya – Fig. S15 and Fig. 3A).

### gProfiler2

Enrichment analysis was performed using *gProfiler2* package [36] (hypergeometric test; multiple comparison correction, ‘gSCS’ method; p-values .01 and .05). Dated variants were subdivided in three time windows (0-300kya, 300kya-500kya and 500kya-1mya) and variant-associated genes (retrieved from [15]) were used as input (all annotated genes for *H. sapiens* in the Ensembl database were used as background). Following [17], variation potential directionality scores were calculated as the sum of all variant effects in a range of 1kb from the TSS. Summary GO figures presented in Figure S12 were prepared with *GO Figure* [59].

For enrichment analysis, the Hallmark curated annotated sets [60] were also consulted, but the dated set of HF variants as a whole did not return any specific enrichment.

## Code URL

https://github.com/AGMAndirko/Temporal-mapping

## Author Contributions

Conceptualization: CB & AA & JM; Methodology: CB & AA & JM; Data Curation: AA & JM; Software: AA & JM; Formal analysis: AA & JM; Visualization: CB & AA & JM & AV & MK & GT; Investigation: CB & AA & JM & AV & MK & GT; Writing – original draft preparation: CB & AA & JM; Writing – review and editing: CB & AA & JM & AV & MK & GT; Supervision: CB; Funding acquisition: CB.

