## Supplementary Figures for "Fine-grained temporal mapping of derived high-frequency variants supports the mosaic nature of the evolution of *Homo sapiens*"

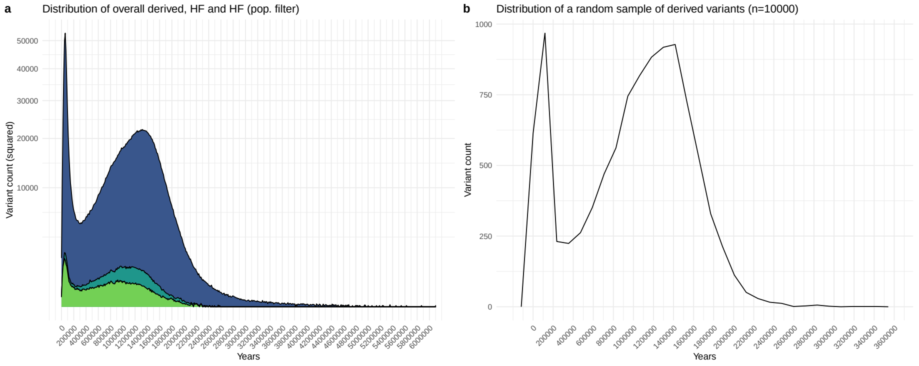


Figure 1: A: Full distribution of derived *Homo sapiens* alleles over time with no frequency cutoff, in HF and the modified population-wise HF subset (see Methods). B: Temporal distribution of a randomly selected sample of derived variants ($n = 10000$).

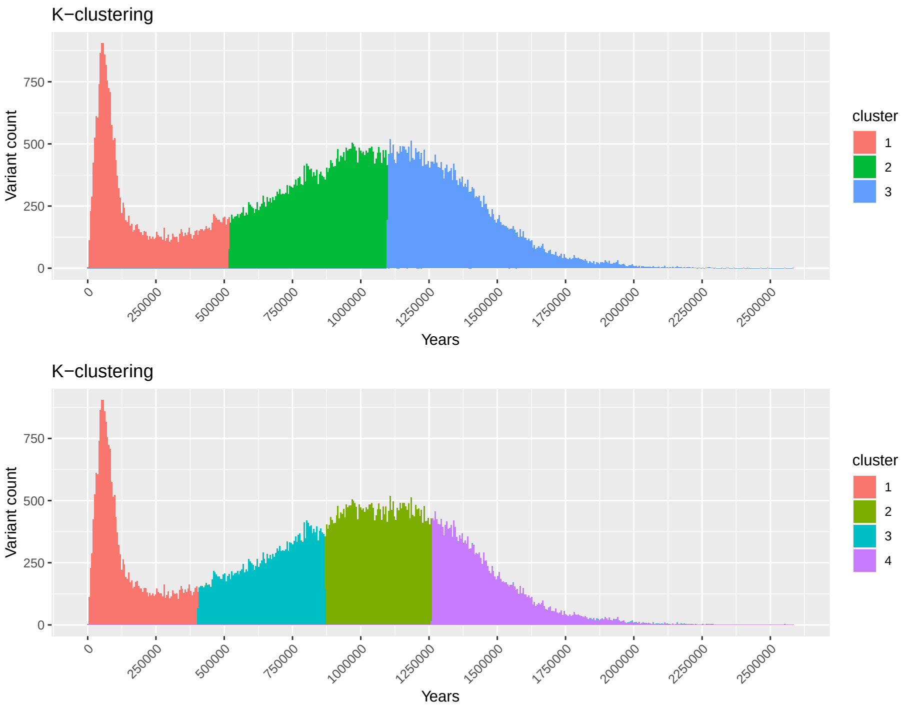


Figure 2: $K$-means clustering analysis of HF variant temporal distributrion.

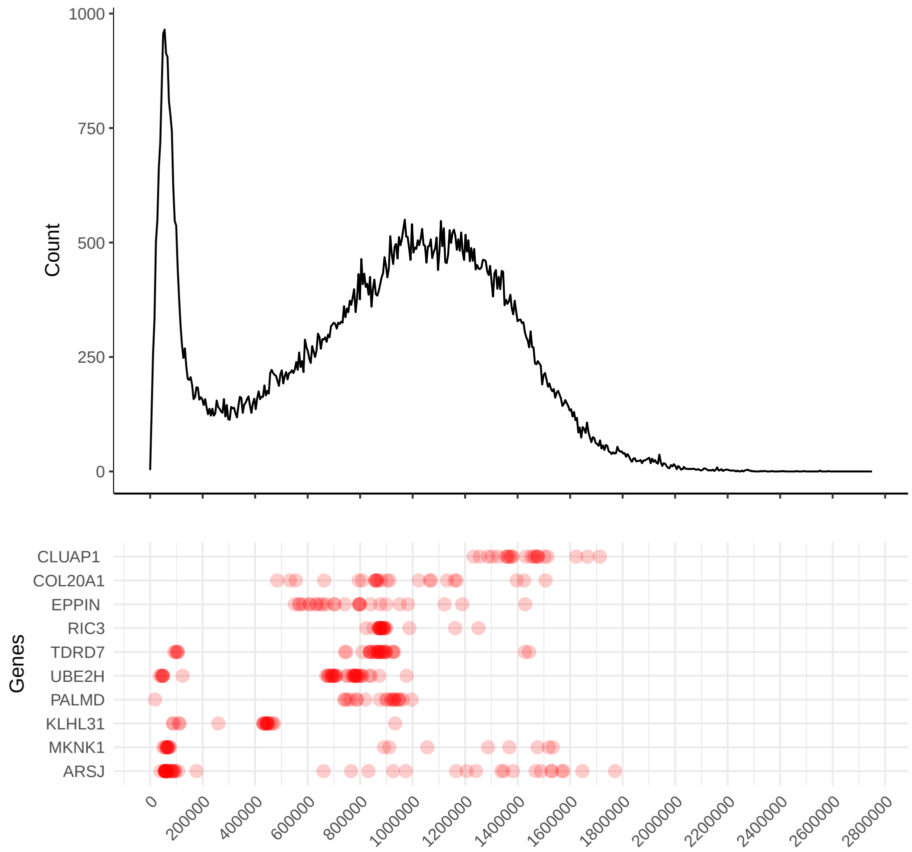


Figure 3: Temporal distribution of variants in genes depleted of archaic-specific variants, as per (1). On top, overall distribution of the HF variant set.

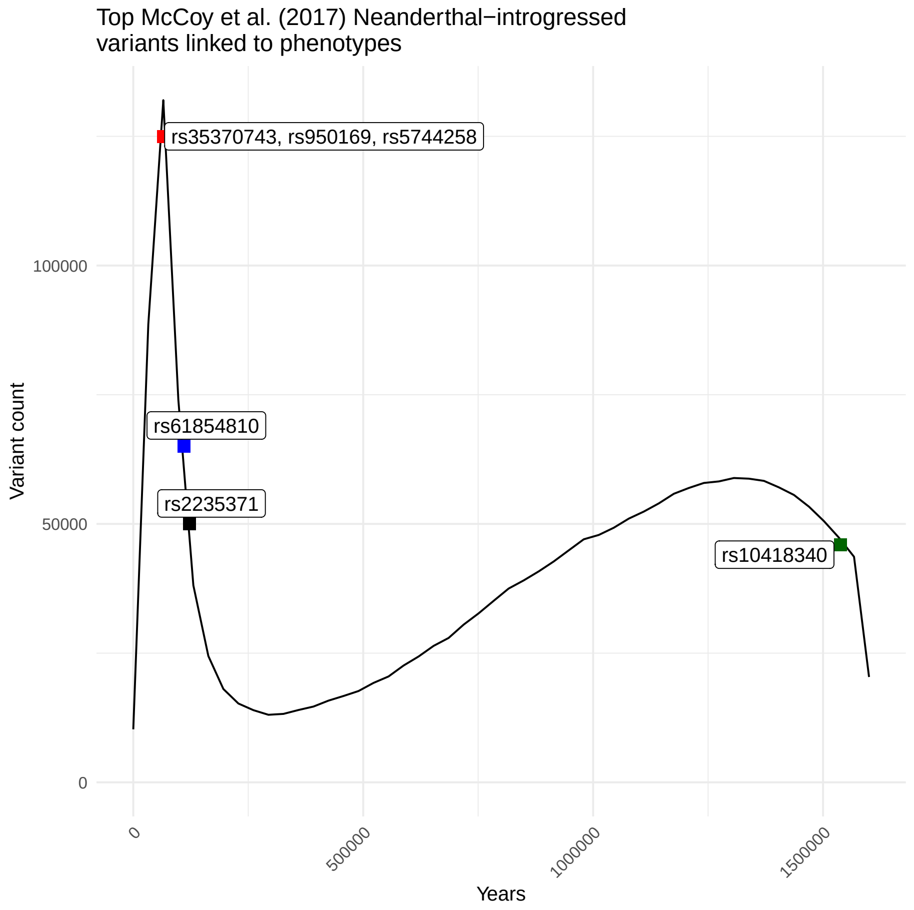


Figure 4: Temporal distribution of introgressed variants linked to phenotypes, as highlighted in Table 1 of (2), compared to the distribution of all derived variants over time.

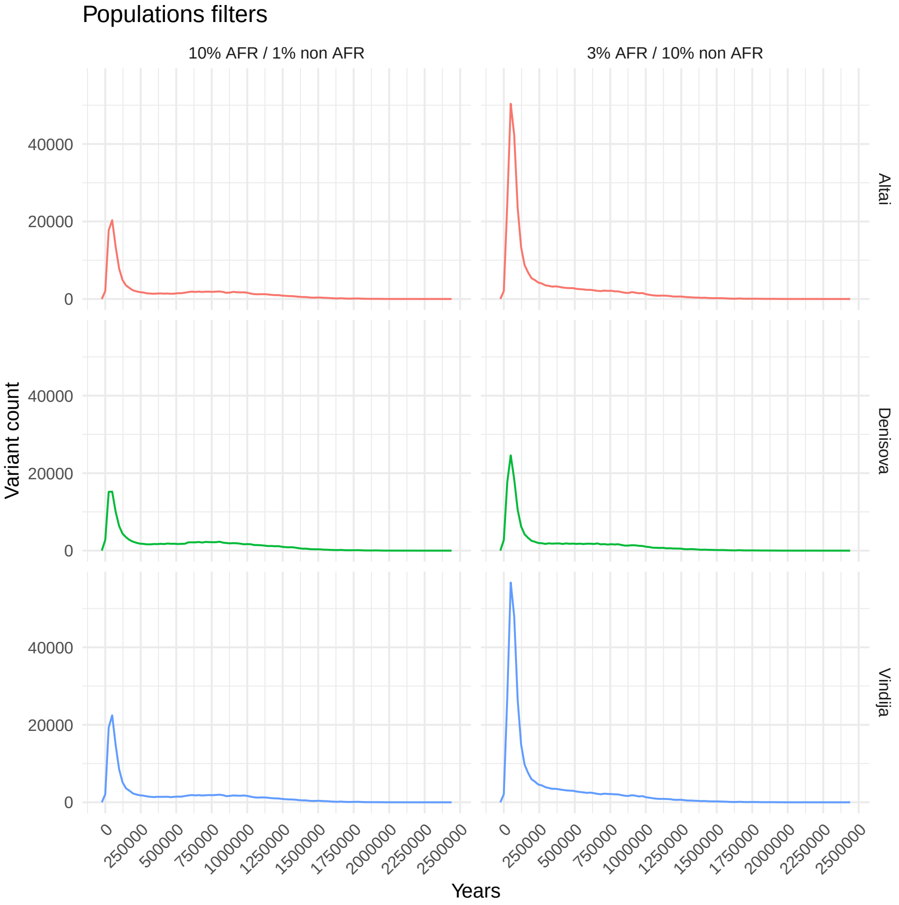


Figure 5: Temporal distribution of variants shared with each of the extinct human genomes after applying specific population frequency filters.These filter include a 10% minor allele frequency cutoff in the African metapopulation (AFR), coupled with a 1% cutoff in the rest of metapopulations, designed to detect potential introgressed alleles brought into the African genetic pool by back-to-Africa migration events. The second filter applied is a 3% cutoff in AFR populations and a 10% threshold in non-African populations, designed to detect the contribution of each extinct human sample to the introgresed variant genetic pool, accounting for a third of that pool to be introduced in AFR populations by back-to-Africa migrations.

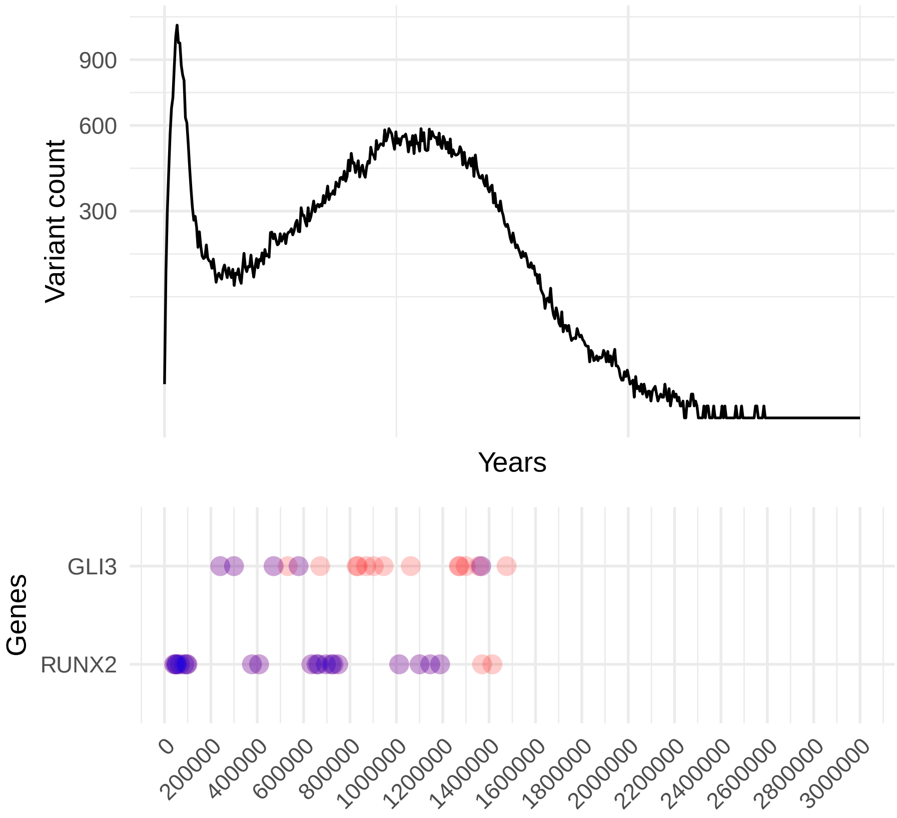


Figure 6: Temporal distribution of HF variants in two genes highlighted in early discussions of selective sweeps: *GLI3* (sweep region from (3) and *RUNX2* (sweep region from (4)). Variants in purple fall within sweep regions.

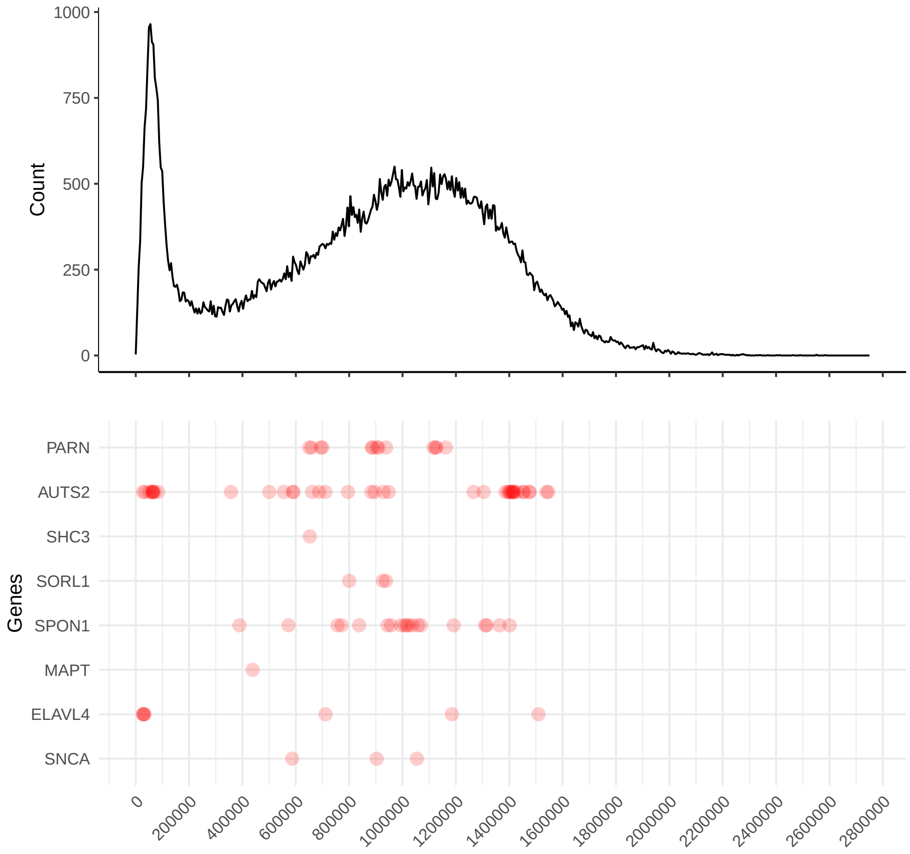


Figure 7: Temporal distribution of variants associated with genes highlighted in (5).

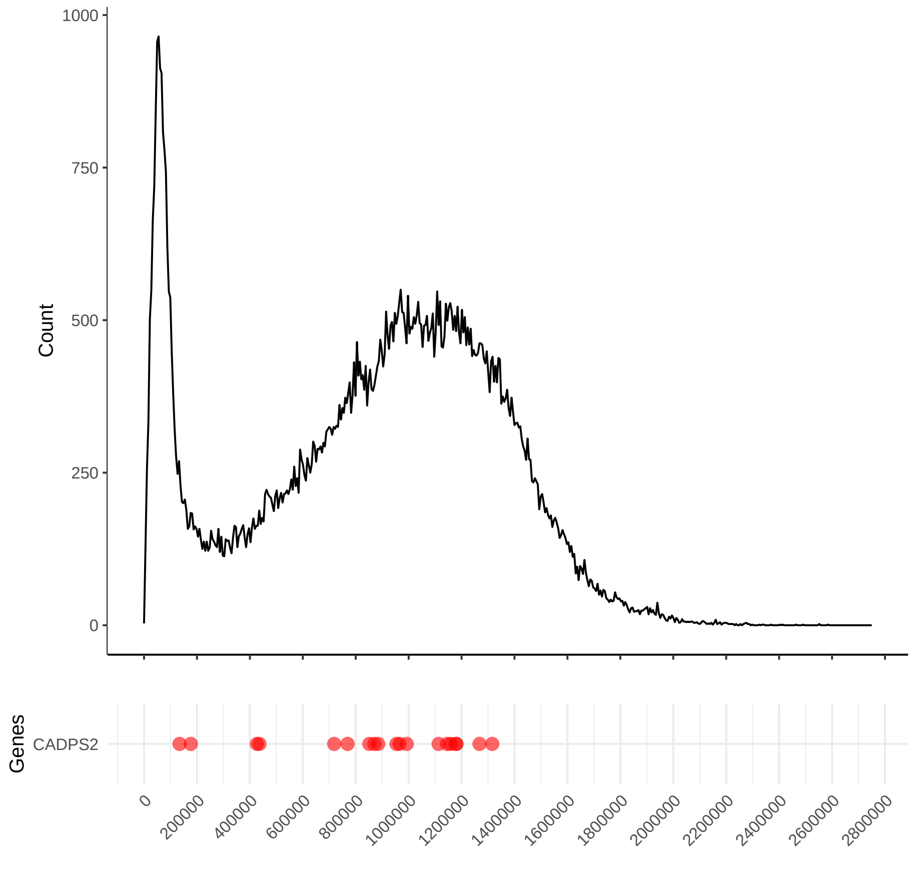


Figure 8: Temporal distribution of variants associated with *CADPS2*. The most recent variants around 200kya in particular capture the reasons this gene was highlighted in (6): "*CADPS2* was identified in (4) as a candidate for selection . . . . The gene has been suggested to be specifically important in the evolution of all modern humans, as it was not found to be selected earlier in great apes or later in particular modern human populations".

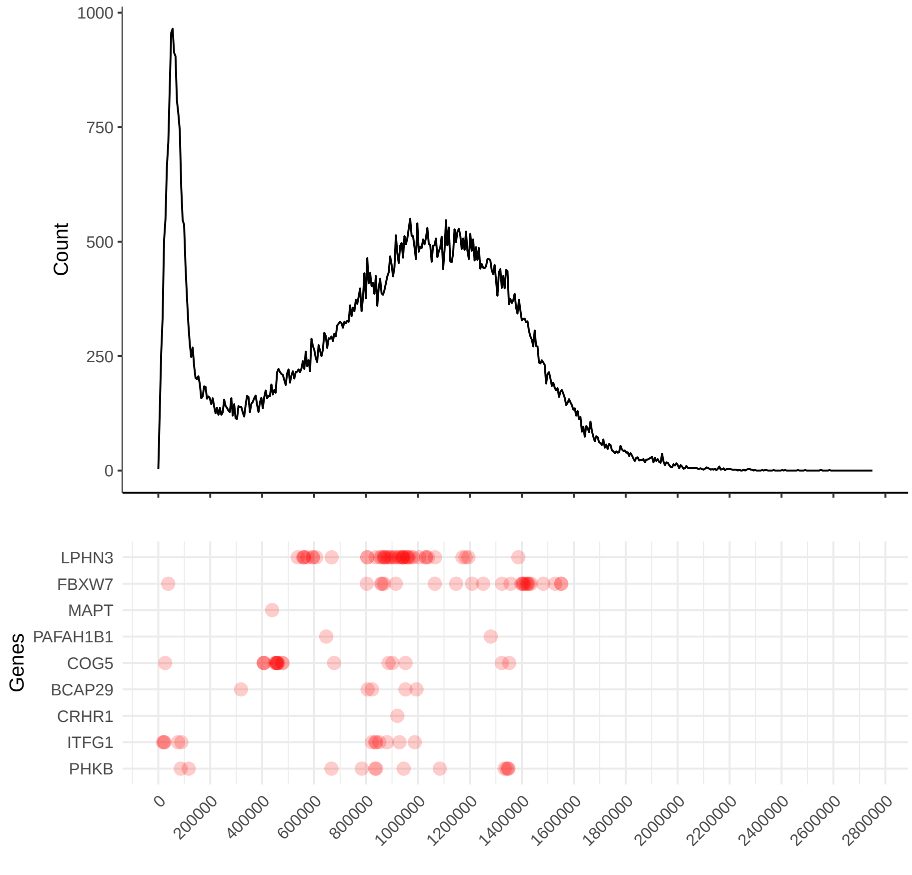


Figure 9: Temporal distribution of variants in genes found in putative positively-selected genetic windows before early *Homo sapiens* population divergence, as per (7). Genes belonging to putative positively selected regions were retrieved from Supplementary Data, section 12 of (7).

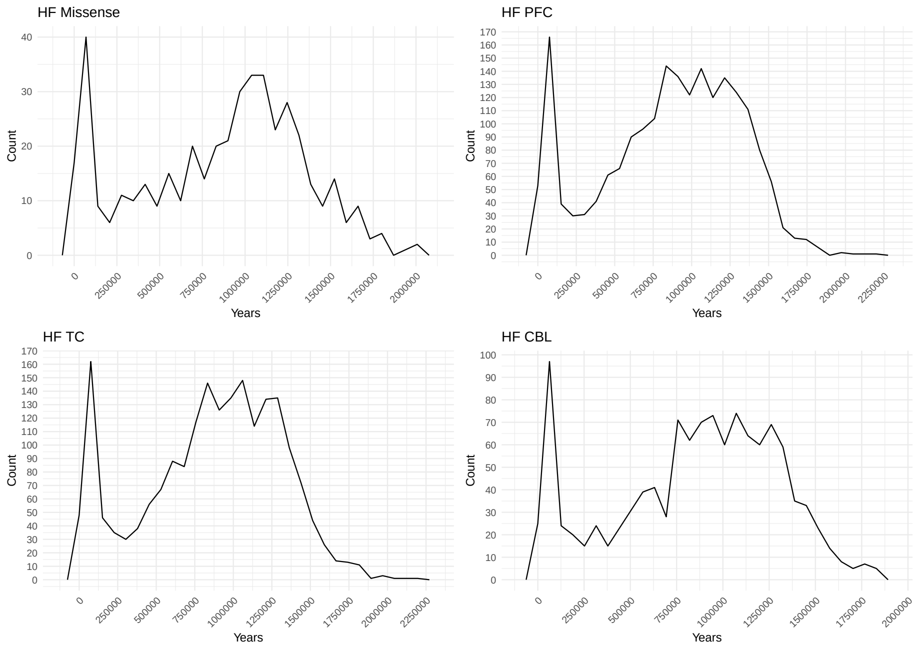


Figure 10: Temporal distribution of high-frequency missense and regulatory variants. Missense variants derived from (1); enhancer annotations for the prefrontal, temporal and cerebellar cortices were retrieved from (8). The difference between the two total maximum counts in the left to the right peak is more pronounced in the cerebellum and prefrontal cortices (23 and 22 more variants mapped to the left maximum peak, respectively). This same difference for missense variants is reduced to only 7 more variants mapped to the left maximum peak. For the temporal cortex, this difference amounts to 14 mapped variants.

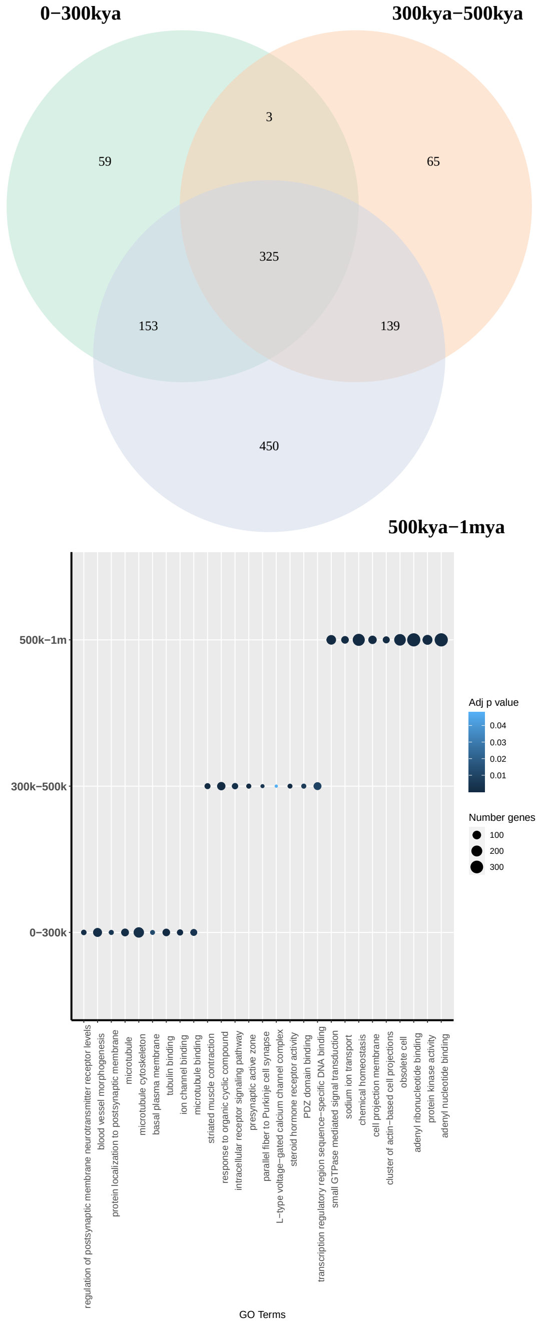


Figure 11: GO terms results when thresholding by an adjusted $p$-value of 0.05. Venn diagram (top) shows number of unique and shared GO terms across periods. Dot plot (bottom) highlights the top 3 GO terms by significance for each period.

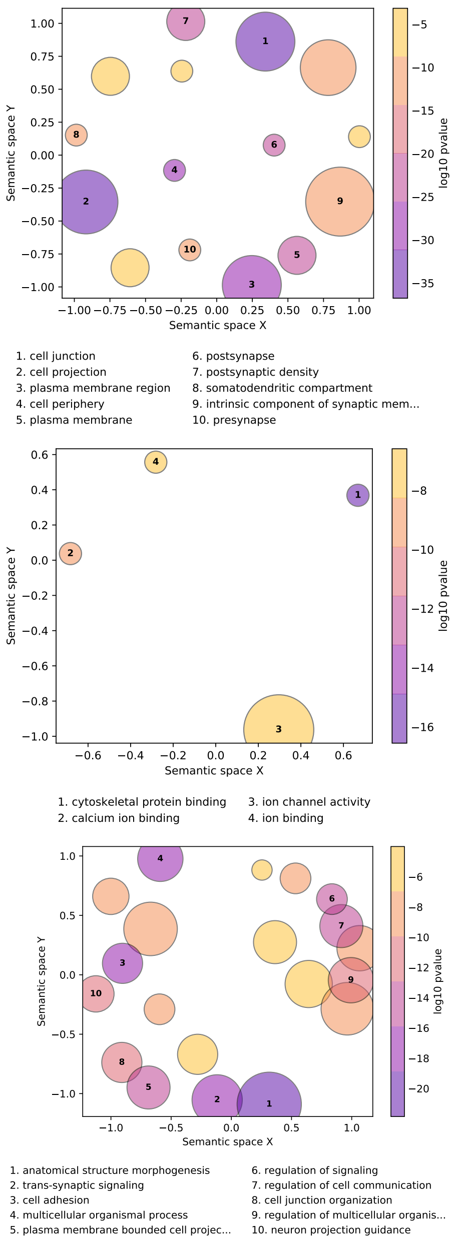


Figure 12: GO term reduction of shared terms across time windows (center of Venn diagram in Fig. 2A)

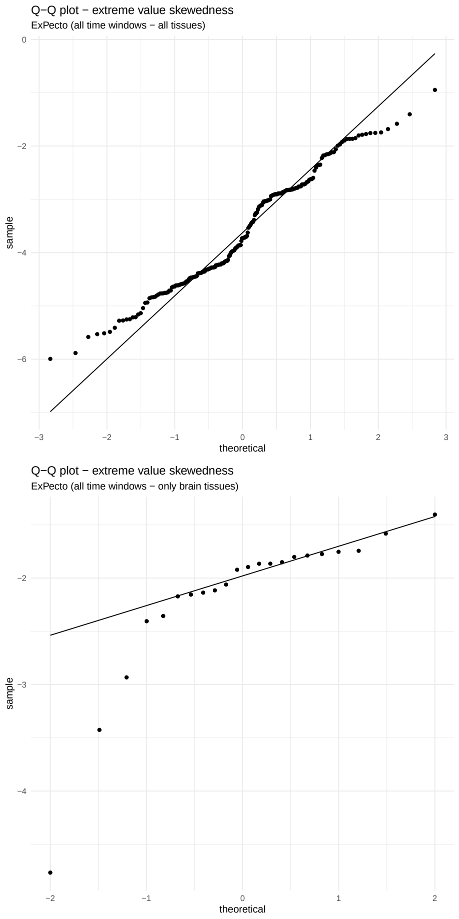


Figure 13: Quantile-quantile plots of predicted expression values skewness for all the tissue models included in (9) and the selected brain-related regions.

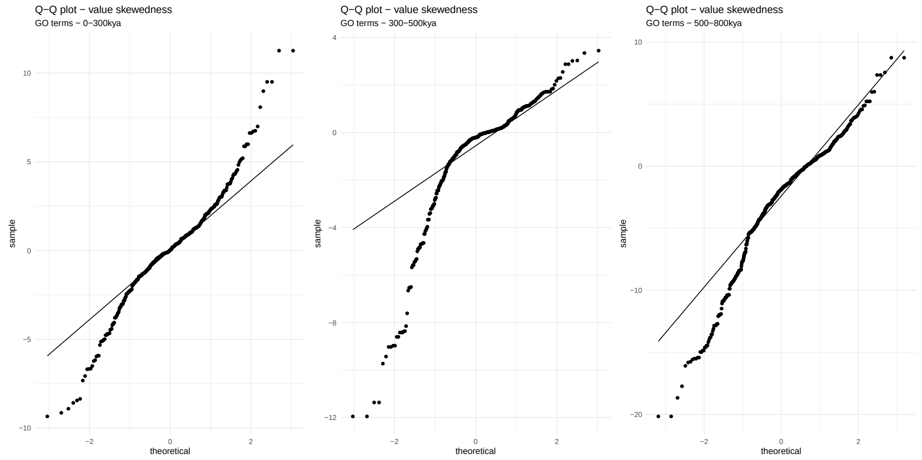


Figure 14: Quantile-quantile plots of predicted expression values of variants associated with GO-enriched genes skewness, divided in three time periods (0-300kya, 300-500kya and 500-800kya). Applied to brain tissue expression only.

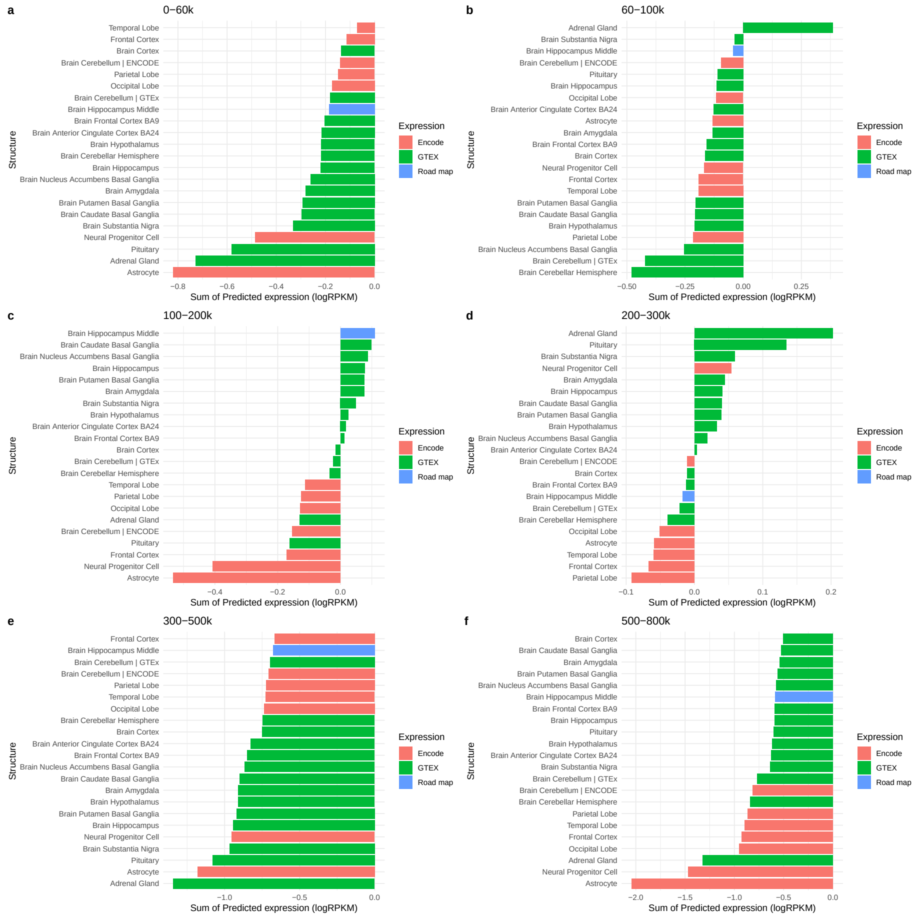

Figure 15: Cumulative predicted expression values by brain-related structure and time window (in years). Color legend indicates data source of prediction models.

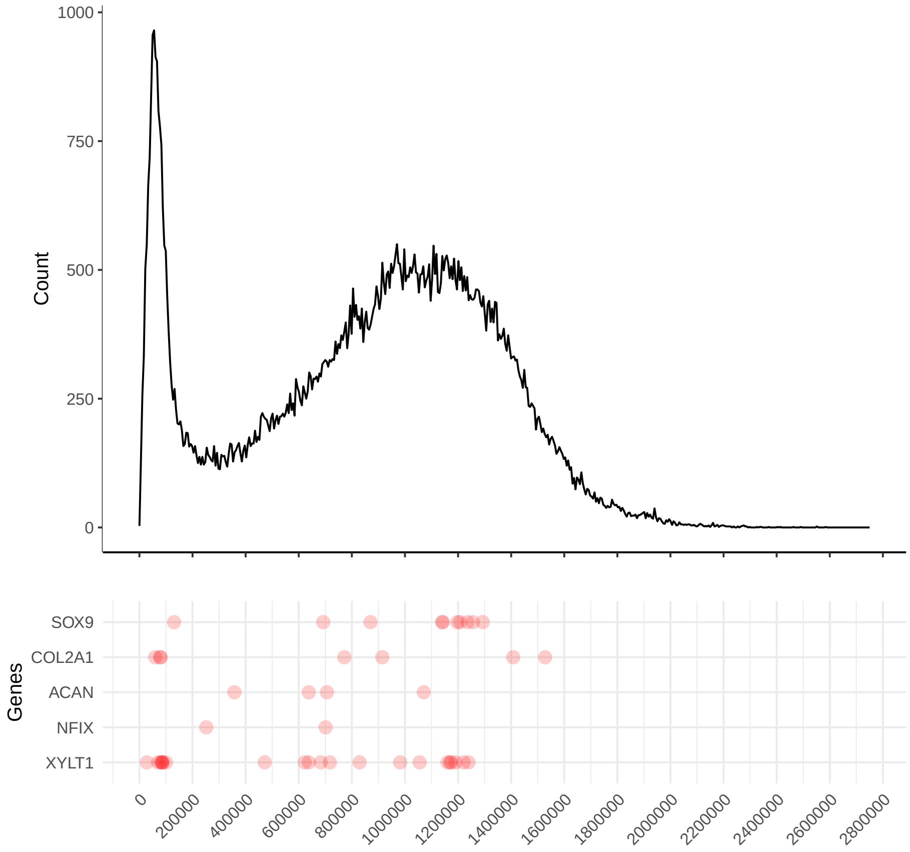


Figure 16: Temporal distribution of variants in genes exhibiting differential methylation profiles in (10) linked to modern human facial traits.

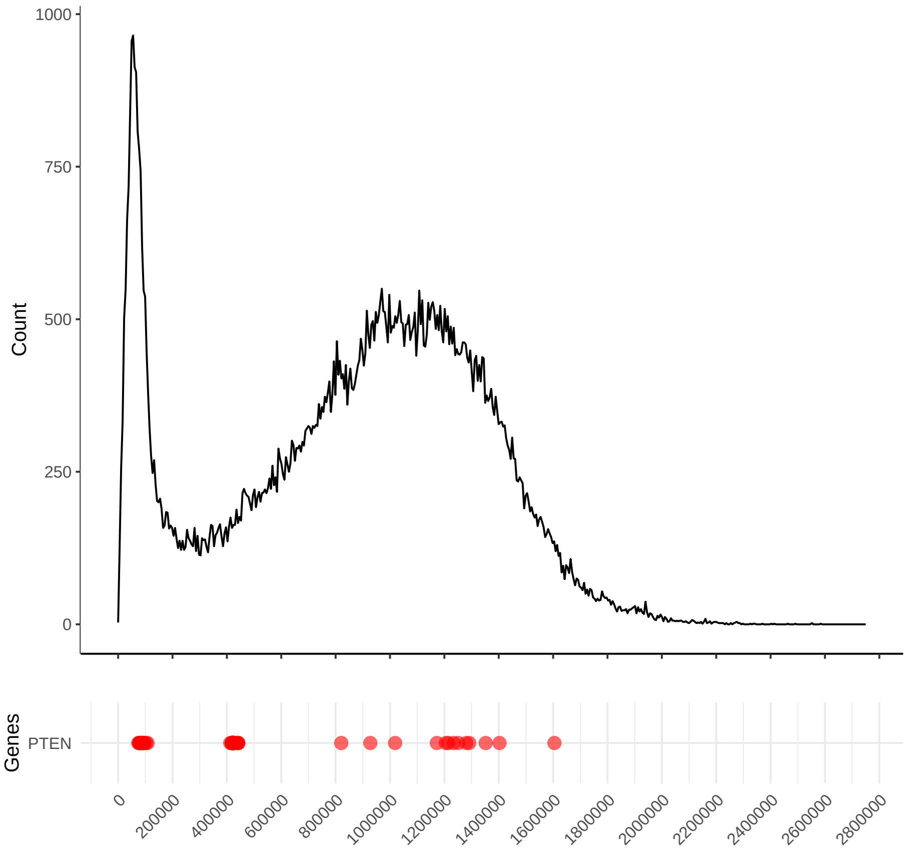


Figure 17: Temporal distribution of variants in $PTEN$, a gene highlighted in (1). The gene displays HF variants clustering around the two periods highlighted in the main text (around 100kya and around 400kya.

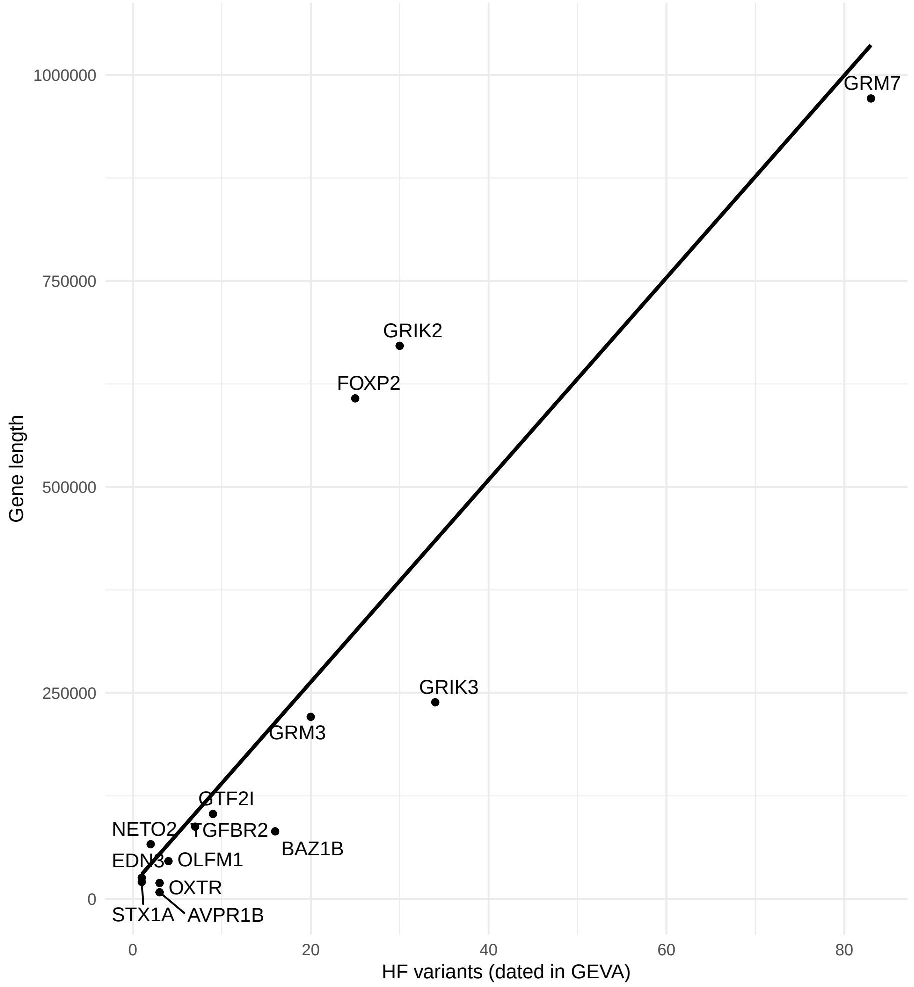


Figure 18: Relationship between gene length and HF variants mapped in GEVA, along with line of best fit.
